# Early development of cerebellar functional connectivity

**DOI:** 10.64898/2026.10.22.760960

**Authors:** Nina Treder, Sunniva Fenn-Moltu, Stuart Oldham, Qianwen Chang, Jonathan O’Muircheartaigh, Dimitar Kostadinov, Tomoki Arichi, Gráinne M. McAlonan, Dafnis Batalle

## Abstract

The cerebellum makes a critical contribution to cognitive and behavioural function, extending well beyond motor control, and is recognised as an important determinant of neurodevelopmental outcome. Despite this, little is known about the foundations of cerebellar functional organisation in early life. Addressing this gap is essential for establishing typical developmental trajectories against which atypical development can be assessed. Here, we analysed fMRI data from term-born neonates (n=155) scanned between 37 and 44 weeks postmenstrual age and mapped intra-cerebellar and cerebro-cerebellar functional connectivity across multiple levels of organisation. We identified 13 intrinsic connectivity networks (ICNs) within the neonatal cerebellum that recapitulated adult subdivisions, with two ICNs showing age-related increases in coverage and strength. Analysis of cerebro-cerebellar connectivity identified 15 sensory, motor and association cerebral clusters, seven increasing connectivity with age. Functional gradient analysis identified a continuous axis from somatomotor to association regions within intra-cerebellar and cerebro-cerebellar organisation, which became more pronounced with age. Together, these findings suggest that the foundations of adult cerebellar functional organisation are present at birth but undergo rapid and regionally heterogeneous maturation over the first weeks of postnatal life. This developmental framework provides a normative reference for understanding cerebellar development and its implications for later neurodevelopmental variation.

## Introduction

The cerebellum is increasingly recognised as a central contributor to human behaviour through its extensive reciprocal connections with the cerebrum (De Benedictis et al., 2022; Ramnani, 2006; Stoodley & Limperopoulos, 2016). Beyond its established role in motor control, converging evidence demonstrates cerebellar involvement in cognitive, affective and executive processes. Consistent with this, differences in cerebellar development have been demonstrated to be an important determinant of neurodevelopmental outcomes (Khan et al., 2015; Sathyanesan et al., 2019; Wang et al., 2014), highlighting the importance of understanding how cerebellar function emerges during early life. Despite growing recognition of the cerebellum’s importance for later cognition and behaviour, little is known about how its functional organisation emerges during the earliest stages of development. This represents a critical gap, as the perinatal period encompasses one of the most dynamic phases of cerebellar maturation, characterised by rapid growth (Holland et al., 2014) and extensive molecular and cellular reorganisation (du Plessis et al., 2018; Garel et al., 2011). Yet, current models of cerebellar functional architecture predominantly rely on studies of older children and adults, requiring assumptions about when and how mature organisational features first emerge during development. Establishing the functional organisation of the cerebellum during the neonatal period is therefore essential for understanding the developmental foundations of mature cerebellar organisation, providing a normative framework against which later developmental variation can be interpreted.

In adults, the cerebellum can be characterised across multiple levels, including discrete functional territories, distributed cerebro-cerebellar networks and more continuous measures of macroscale organisation. The anterior lobe (lobules I-V) is primarily associated with somatomotor control and proprioception (Stoodley & Schmahmann, 2009) and receives input from cerebral somatomotor regions (Kelly & Strick, 2003), whereas the posterior lobe (lobules VI-XI, including Crus I/II) predominantly receives input from the prefrontal and parietal cortex and supports affective and higher-order cognitive processes (Buckner et al., 2011; King et al., 2019; Manni & Petrosini, 2004; Salmi et al., 2010; Stoodley & Schmahmann, 2009). Beyond theses discrete functional regions in the cerebellum, recent studies have demonstrated that the organisation of its connectivity can also be described along continuous functional gradients, which capture smooth transitions in connectivity rather than abrupt boundaries between specific regions (Guell et al., 2018). These gradients extend from unimodal somatomotor to transmodal association regions, mirroring the canonical macroscale organisation observed across the cerebral cortex (Dong et al., 2022; Guell et al., 2018; Margulies et al., 2016). Together, these organisational features provide complementary descriptions of mature cerebellar function, yet it remains unclear when and how they emerge during early development.

Initial evidence suggests that aspects of adult-like cerebellar organisation may already be present early in life. ROI-based studies have identified selective cerebro-cerebellar connectivity patterns in neonates (Herzmann et al., 2019), while more recent work spanning infancy and early childhood has reported emerging cerebellar resting-state networks and patterns of cerebro-cerebellar connectivity (Lyu et al., 2025; Tikoo et al., 2025). However, these studies have primarily focused on predefined regions of interest, broad developmental periods extending across years rather than the earliest stages of postnatal refinement, or were conducted at lower spatial resolution, potentially limiting characterisation of neonatal cerebellar organisation. As a result, current evidence provides only a partial description of early cerebellar function with little insight into how intrinsic networks, cerebro-cerebellar connectivity patterns, and macroscale organisational gradients are structured and refined during the weeks immediately following birth.

Here, we provide a comprehensive characterisation of early cerebellar functional organisation in a cohort of term-born neonates and define, for the first time, the principal axes of neonatal intra-cerebellar and cerebro-cerebellar functional connectivity using gradient analysis. First, we define data-driven intrinsic connectivity networks (ICNs) within the neonatal cerebellum and examine age-related variation in their connectivity strength and spatial coverage across the first weeks of postnatal life. Second, we map voxel-wise cerebro-cerebellar functional connectivity in the cerebrum and cerebellum, characterise its organisational structure, and assess developmental variation in the early neonatal period. Third, we apply gradient analysis to characterise the macroscale organisation of neonatal cerebellar and cerebro-cerebellar connectivity, providing insight into whether canonical and continuous principles of adult cerebellar organisation are already evident during early postnatal development.

## Results

### Sample

After extensive qualitative and quantitative assessment of cerebellar fMRI signal quality (see Methods, and Supplementary Figure S1), exclusion of datasets with incomplete cerebellar coverage, uncorrectable signal artefact due to excessive head motion, a history of preterm birth and brain injury, we included 155 (of 809) fMRI datasets acquired from term-born neonates (67 females; gestational age (GA) at birth > 37 weeks) acquired as part of the developing Human Connectome Project (dHCP) (release 3, Edwards et al., 2022). Average GA at birth was 40.1 (SD = 1.25) weeks and the average postmenstrual age (PMA) at scan was 41.4 (SD = 1.66) weeks. The sample was split into two subsets based on specific analytical goals. Subset 1 data (20 term-born neonates; GA at birth: 38-42 weeks; PMA at scan: 43-44 weeks) were used to identify group-level functional networks, connectivity and gradients, that were then later used to compare neonates at the individual level. Subset 2 data (135 term-born neonates; GA at birth, 37-42 weeks; PMA at scan, 37-44 weeks) were used to characterise variations within functional connectivity features across age at the individual level, and to chart early functional development of the cerebellum (Supplementary Figure S1).

**Table 1.** Subject demographics and follow-up at 18 months. Mean, standard deviation (SD) and range displayed for each continuous variable. Number and percentages displayed for categorical variables.

| **Subset Analysis** | **Subset 1 Group-level** | **Subset 2 Subject-level** | **Test statistic** | **p-value** |
| --- | --- | --- | --- | --- |
| **Number of participants** | 20 | 135 |  |  |
| **Sex (N females) [%]** | 9 [45%] | 58 [43%] | 0.030<sup>a</sup> | $p = 0.864$ |
| **GA at birth in weeks [SD] (range)** | 40.57 [0.95] (38-42) | 40.01 [1.28] (37-42) | 1.828<sup>b</sup> | $p = 0.068$ |
| **PMA at scan in weeks [SD] (range)** | 43.42 [0.58] (43-44) | 41.09 [1.56] (37-44) | 6.048<sup>b</sup> | $p < 0.001$ |
| **% FD outliers [SD] (range)** | 3.80 [3.11] (0.61-9.70) | 5.58 [2.62] (0.26-10.00) | -2.597<sup>b</sup> | $p = 0.01$ |

<sup>a</sup> $\chi^2$ -Test <sup>b</sup> Z (Mann Whitney U-Test)

### Cerebellar intrinsic connectivity networks are present at birth and mature during the neonatal period

To identify spatially independent patterns of correlated functional activity within the cerebellum, we applied spatial Group-ICA (Beckmann C Smith, 2004) to fMRI data in subset 1 from term-born infants scanned between 42.5-44.5 weeks PMA (*n = 20*) (Subset 1) (Figure 1A). We identified 12 unique ICNs located within the cerebellar cortex and one ICN corresponding to the dentate nucleus of the cerebellum. The ICNs were distributed across Crus I/II, lobules V-VIII, and the vermis, revealing a diverse set of early functional networks spanning both sensorimotor-related and association related regions (Figure 2A).

**Figure 1.**
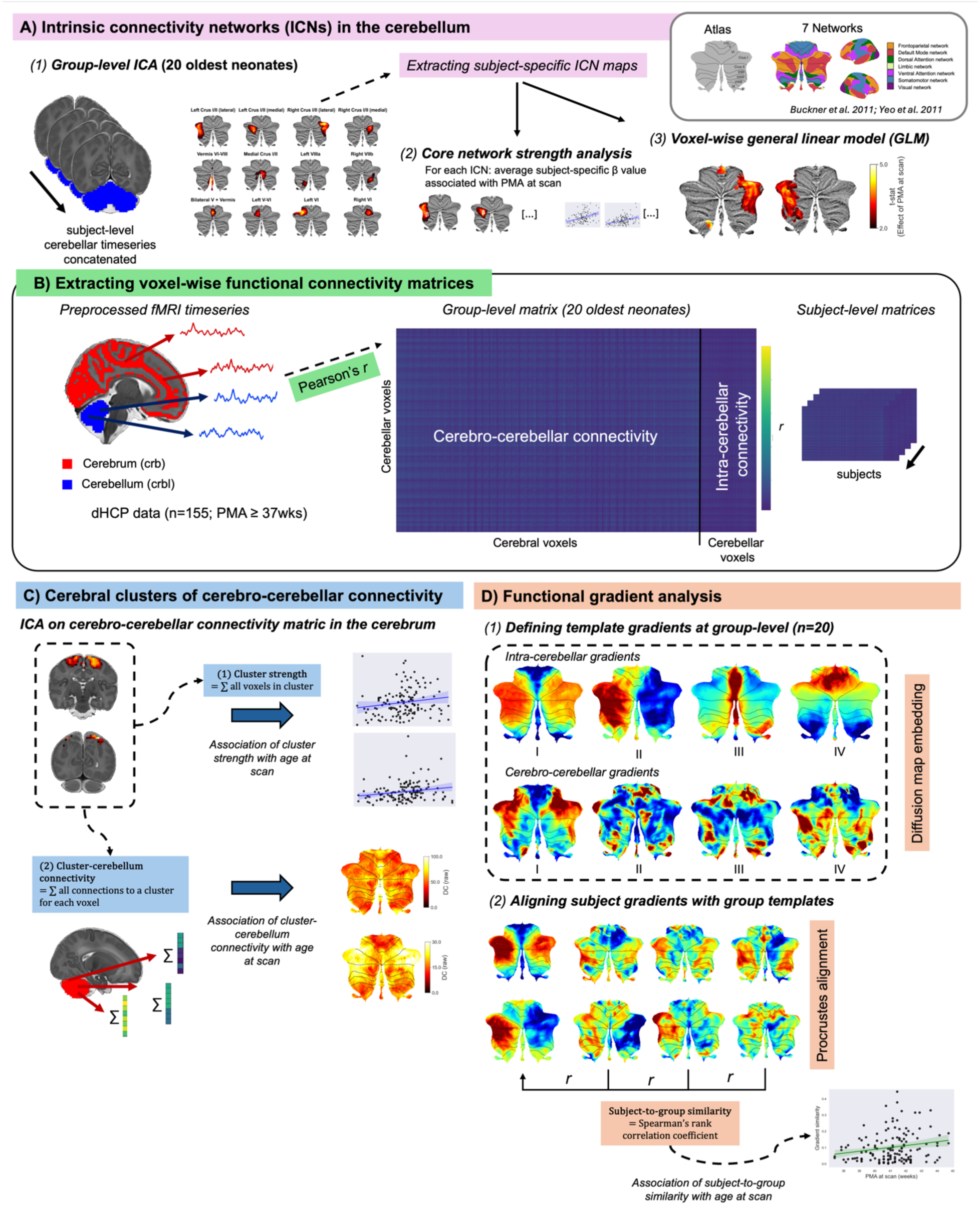
Summary of methodology. A) Intra-cerebellar intrinsic connectivity networks (ICNs) derived with group-ICA. Subject-specific ICNs were extracted using dual regression and core network strength per ICN was quantified by averaging subject-specific β values. Spatial ICN structure and core network strength was associated with postmenstrual age (PMA) at scan. For anatomical reference, cerebellar anatomy and functional topography in the cerebellum derived from adults (Buckner et al., 2011) based on the Yeo-7 networks (Yeo et al., 2011) is displayed on cerebellar flatmaps (top right) B) Pre-processed neonatal fMRI data were selected from the dHCP dataset and fMRI time series in the cerebrum (red) and cerebellum (blue) were extracted for all subjects. Voxel-wise functional connectivity matrices were computed for each subject using Pearson’s correlation coefficient. The group connectivity matrix was computed from the 20 oldest neonates using Fisher’s r-to-z transformation. C) Clusters of cerebro-cerebellar connectivity in the cerebrum were derived using ICA at the group-level. Cluster strength and cluster-cerebellum connectivity were associated with PMA at scan. D) Intra-cerebellar (top) and cerebro-cerebellar (bottom) functional gradients in the neonatal cerebellum were defined using diffusion map embedding on the group connectivity matrix and projected onto cerebellar surfaces. Subject-level gradients were derived and aligned with group templates using Procrustes alignment. Subject-level gradients were then correlated with group templates using Spearman’s rank correlation coefficient to obtain subject-to group gradient similarity.

**Figure 2.**
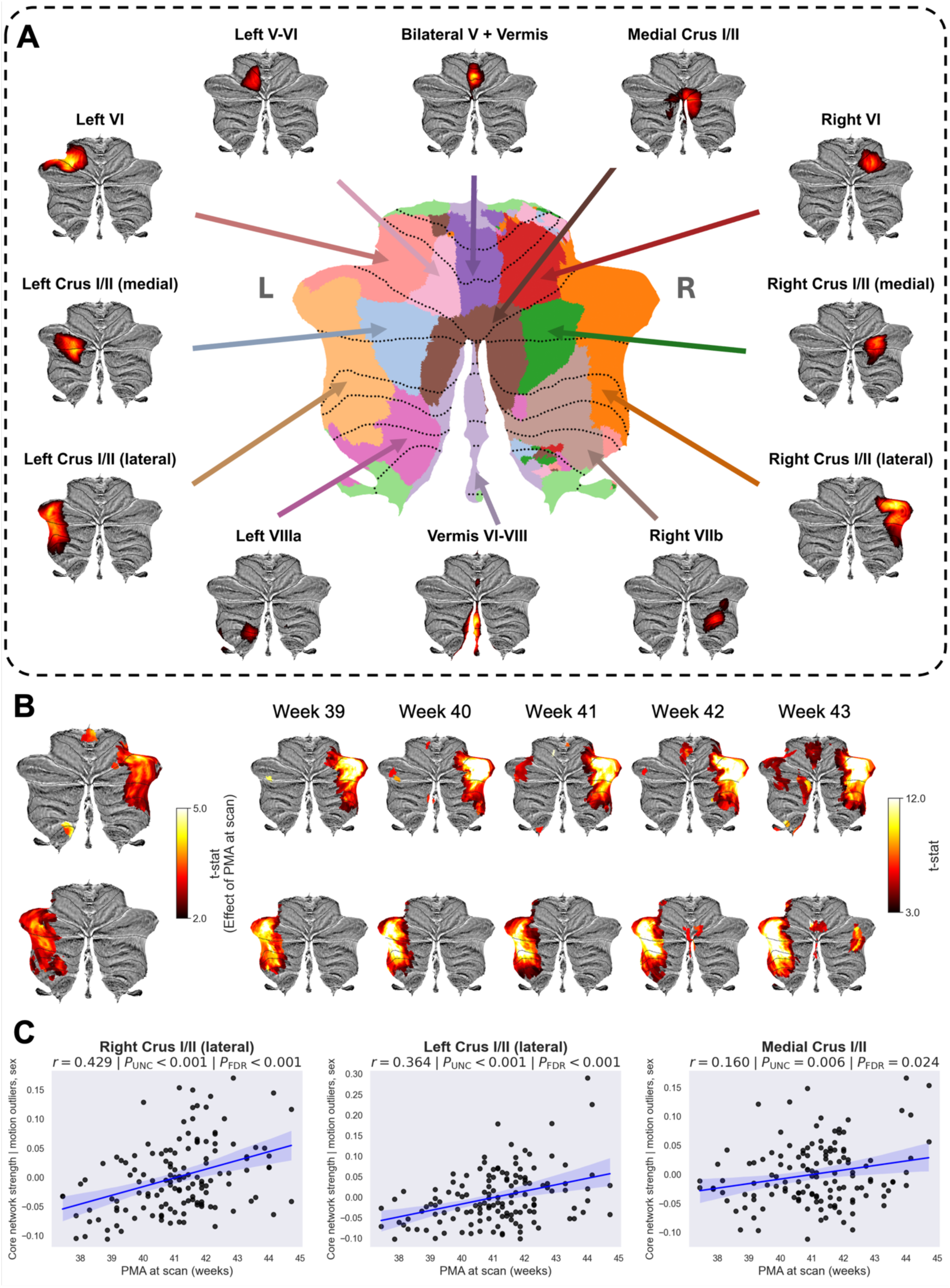
Intrinsic connectivity networks (ICNs) in the neonatal cerebellum. (A) Group-level cerebellar ICNs displayed with anatomical labels on separate flatmaps and as one cerebellar parcellation in the center. Group-ICA results are displayed thresholded at Z > 3 at surface-level with left (L)-right (R) orientation as indicated. (B) Changes in ICN coverage with age and weekly maturation in functional network structure of right lateral Crus I/II (top) and left lateral Crus I/II (bottom). (C) Association of core network strength of ICNs and PMA at scan. Scatterplots display mean within-network β residuals adjusted for sex and motion outliers of cerebellar ICNs in the right lateral Crus I/II, left lateral Crus I/II and medial Crus I/II. Regression line in blue, with shaded areas indicating 95% confidence intervals.

Age-related maturation of intra-cerebellar functional connectivity was assessed in subjects of subset 2, aged 37 to 44 weeks PMA at scan, by modelling the effect of PMA at scan on subject-specific ICN coverage and amplitude. Subject-specific spatial maps for each ICN were estimated using dual regression (Nickerson et al., 2017) and regressed against PMA at scan, controlling for sex and motion. Voxel-wise inference was performed using non-parametric permutation testing implemented in FSL randomise (10,000 permutations) with threshold-free cluster enhancement (TFCE) (Winkler et al., 2014) and family-wise error correction. Two ICNs in the left and right Crus I/II showed increasing connectivity with PMA at scan. Developmental changes in network structure were also qualitatively characterised by categorising term-born infants into weekly groups and generating group-specific spatial maps. Both left and right Crus I/II networks exhibited a clear increase in amplitude and spatial coverage across the early postnatal period, consistent with ongoing maturation of higher-order cerebellar regions (2). There were no ICNs showing a negative association with PMA at scan (Figure 2B).

In order to summarise connectivity strength for each ICN, core network strength was calculated as the mean β-parameter value in the ICN-specific spatial map of each subject (obtained from stage two of dual regression) after masking by the corresponding group-ICA network template thresholded at Z > 3 (Eyre et al., 2021). This allowed for the measurement of within-network functional connectivity by testing associations of core network strength with PMA at scan using partial correlations controlling for sex and motion (% FD outliers). Three ICNs showed a positive association between core network strength and age, namely ICN in the right Crus I/II (r = 0.457, p_FDR_ < 0.001) and left Crus I/II (r = 0.435, p_FDR_ < 0.001), as well as the ICN in the Medial Crus I/II region (r = 0.263, p_FDR_ = 0.024) (after False Discovery Rate [FDR]-correction (Figure 2C).

### Cerebro-cerebellar connectivity reinforces cerebral motor and association network structure

We first constructed voxel-wise functional connectivity matrices between the cerebrum and the cerebellum using Pearson’s correlation coefficients in Subset 1 at the group-level (Figure 1B). Cerebral voxels were then clustered according to their correlation profiles with cerebellar voxels using ICA (dimensionality = 20) (Figure 1C). 15 clusters across motor, association and visual regions of the cerebrum were retained based on visual ckassification and a quantitative cluster-size threshold derived from a null distribution generated using 1,000 permutations (Supplementary Figure S2). We quantified the strength of cerebro-cerebellar connectivity by summing for each subject, all cerebellar connections with voxels within each cluster (thresholded at Z ≥ 3), yielding a single cluster-strength value per cluster. We then modelled the association between PMA at scan and cluster strength using partial correlations, controlling for sex and motion. Seven cerebral clusters increased in strength with PMA at scan (Figure 3A), including the visual association (left) (r=0.204, p_FDR_=0.039), posterior parietal (r=0.279, p_FDR_=0.009), medial visual (r=0.385, p_FDR_< 0.001), somatomotor (r=0.230, p_FDR_=0.039), supplementary motor I (r=0.204, p_FDR_=0.039), supplementary motor II (r=0.210, p_FDR_=0.039) and precuneus (left) (r=0.204, p_FDR_=0.039) cluster (Figure 3B).

**Figure 3.**
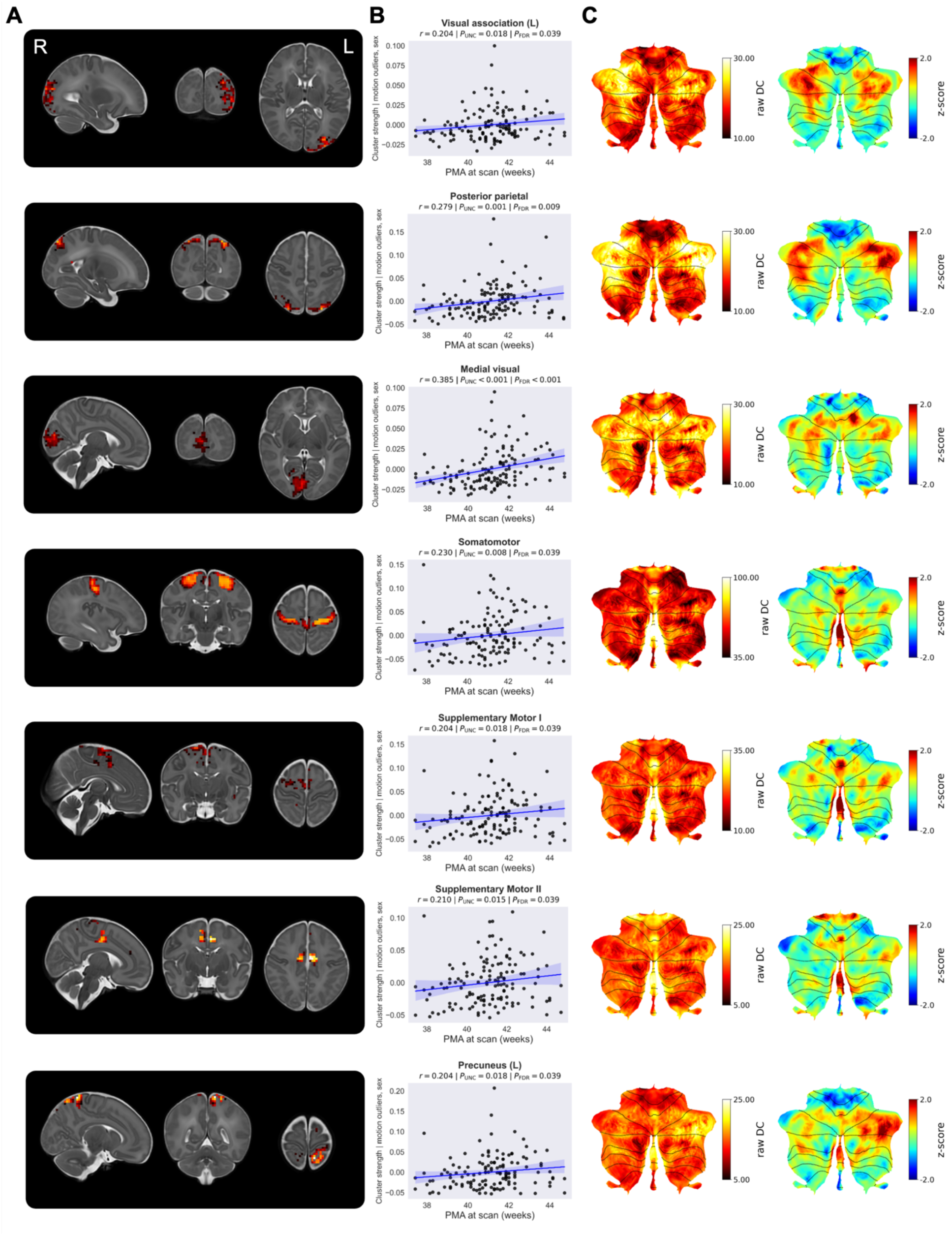
Cerebral clusters underlying cerebro-cerebellar connectivity. (A) Group-level clusters, identified using ICA (dimensionality = 20), showing significant increase in cluster strength with PMA at scan, thresholded at Z ≥ 3. (B) Association between cluster strength and PMA at scan. Scatterplots show cluster strength residuals after adjusting for sex and motion outliers. Regression line in blue indicates linear regression fit with shaded 95% confidence intervals. (C) Raw (left) and z-scored (right) degree centrality (DC) maps of cerebellar connectivity with the respective cerebral clusters, displayed on the cerebellar surface.

To identify which cerebellar regions were most strongly connected to each cluster, we summed for each cerebellar voxel at the group-level connectivity with voxels in each cerebral cluster separately, resulting in cluster degree centrality (DC) maps. To facilitate visual interpretation and reference across clusters, we additionally generated z-scored cluster DC maps, which normalise connectivity values relative to the whole-cerebellum distribution of cluster DC values and highlight regions showing relatively stronger or weaker connectivity with a specific cluster. Visual-related clusters (visual association [L] and medial visual) and association-related clusters (posterior parietal, precuneus [L]) show strongest connectivity with the Crus I/II region of the cerebellar posterior lobe. In contrast, motor-related clusters (somatomotor, premotor I and premotor II) were most strongly connected with the vermis and anterior lobe regions associated with motor coordination (Figure 3C).

Together these findings indicate that cerebro-cerebellar connectivity is already organised along sensory, associative and motor axes in the neonatal period, with modest but significant age-related increases in coupling strength observed across 37-44.5 weeks PMA.

### Intra-cerebellar and cerebro-cerebellar functional gradients are present at birth

Gradient analysis can be used to represent the dominant axes of variance in functional connectivity without requiring a priori network parcellations and has previously been used to identify a principal unimodal-to-multimodal axis of organisation in the adult cerebral cortex and cerebellum (Margulies et al., 2016). To better understand the macroscale functional organisation of both intra-cerebellar and cerebro-cerebellar connectivity, we applied diffusion map embedding to represent the high-dimensional connectivity data in a low-dimensional gradient space (Figure 1D). Functional gradients capture the dominant axes of variation in connectivity, positioning voxels along continuous spatial dimensions according to the similarity of their connectivity profiles (Bernhardt et al., 2022). For both intra-cerebellar and cerebro-cerebellar connectivity, the first four gradients explained approximately 70% of the variance (71.3% and 70.8%, respectively). The principal intra-cerebellar gradient *I* spanned a transmodal-to-unimodal axis, reflecting a topological continuum of organisation from somatomotor regions through attention networks to higher-order regions associated with the cerebral default mode network, similar to previously described gradients in the adult cerebellum (Figure 4A-B). Intra-cerebellar gradient *II* reflected hemispheric differentiation, gradient *III* captured a posterior-anterior lobe gradient, while gradient *IV* captured a transition from the vermis to the cerebellar hemispheres (Figure 4B). Gradients derived from cerebro-cerebellar connectivity yielded a similar transmodal-unimodal gradient *I*, which was strongly correlated with intra-cerebellar gradient *I* (r = 0.718) and *III* (r = 0.512). Gradients *II*-*IV* captured a posterior-anterior axis, indicating partially shared organisational principles across intra-cerebellar and cerebro-cerebellar functional connectivity. (Figure 4C). Higher-order cerebro-cerebellar gradients *II* and *III* showed minimal associations with intra-cerebellar gradients *I-IV*, while cerebro-cerebellar gradient *IV* showed low to moderate correlations with intra-cerebellar gradient *III* (r = 0.285) and *IV* (r = 0.352) (Supplementary Figure S3).

**Figure 4.**
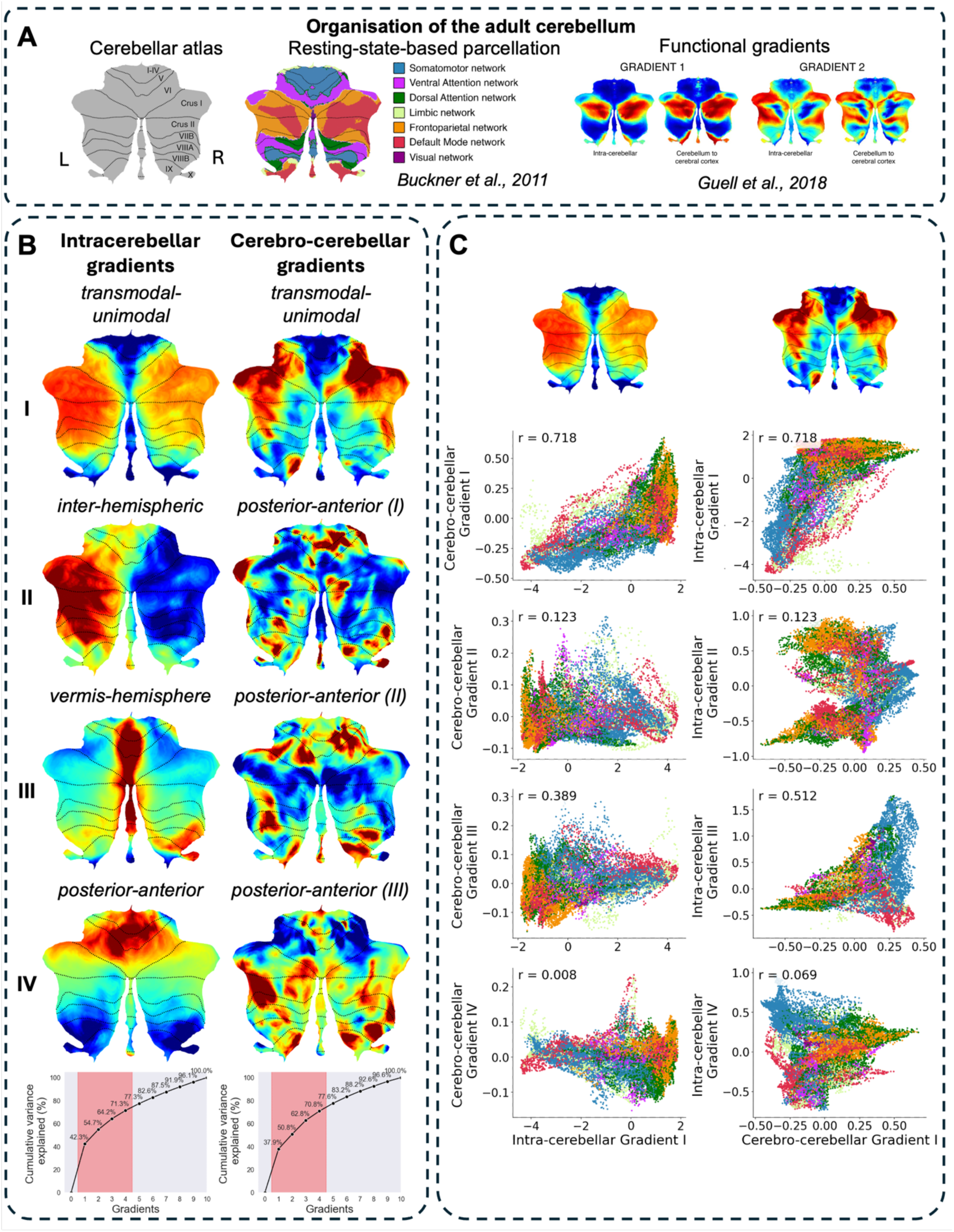
Functional gradients of the neonatal cerebellum. (A) Organisation of the adult cerebellum, including anatomy and topography and resting-state networks shown on cerebellar flatmaps (left) (adapted from the data provided in Buckner et al., 2011); and adult intra-cerebellar and cerebro-cerebellar functional gradients on cerebellar flatmaps (right) (reproduced from Guell et al., 2018, under the terms of the Attribution License CC BY 4.0. (B) Intra-cerebellar (left) and cerebro-cerebellar (right) functional gradients *I*-*IV* represented in adult surface space with cumulative variance explained (%) shown at the bottom. The shaded red indicates selected gradients. (C) Scatterplots (left) show intra-cerebellar gradient *I* plotted against cerebro-cerebellar gradients *I*-*IV*, and cerebro-cerebellar gradient I plotted against intra-cerebellar gradients *I-IV*, with each point representing a cerebellar vertex coloured by resting-state network (from Buckner et al. 2011), and positions determined by gradient values.

### Cerebellar functional gradients represent distinct macroscale network organisation

The relationships between intra-cerebellar and cerebro-cerebellar sub-networks were characterised by assessing sub-network separation in gradient space (Figure 5). In intra-cerebellar gradient space, gradients *I* and *II* showed the strongest sub-network dissociation, i.e., the clearest functional separation. By contrast, comparisons between gradient *I* and gradients *III* and *IV* (Figure 5A), as well as among gradients *II, III* and *IV* themselves (Supplementary Figure S4A), showed greater sub-network intermixing and less distinct functional boundaries, suggesting more fine-grained or overlapping functional distinctions. In cerebro-cerebellar gradient space, gradient *I* similarly showed stronger sub-network separation (Figure 5B) relative to gradients *II, III* and *IV*, which displayed greater functional overlap (Supplementary Figure S4B). Median gradient values for each resting-state network across both intra-cerebellar and cerebro-cerebellar gradient *I* showed that frontoparietal, default mode, and dorsal attention networks were closely located to each other in gradient space, while regions associated with the ventral attention, visual, somatomotor and limbic networks formed a separate grouping (Supplementary Figure S5).

**Figure 5.**
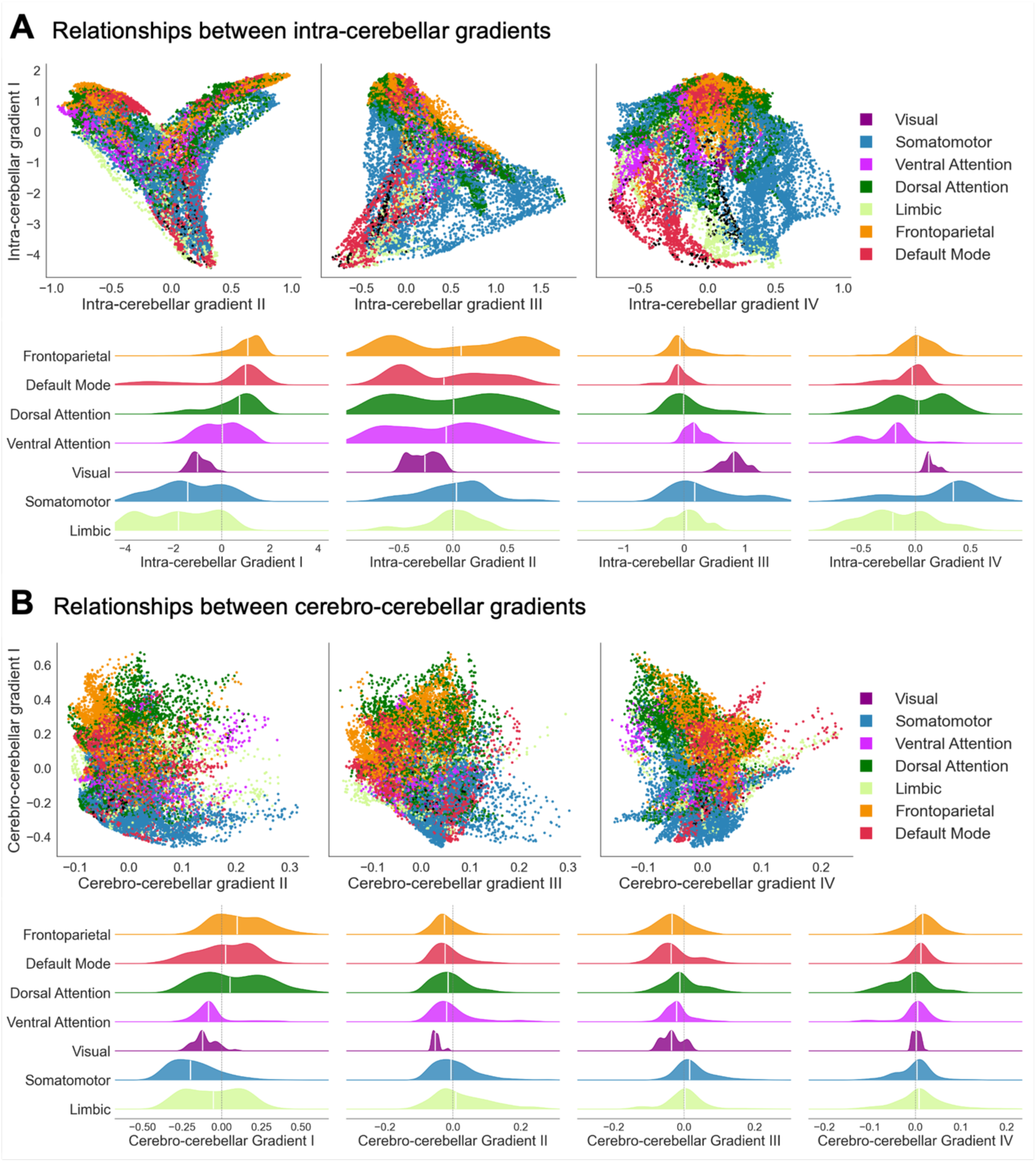
Relationships between cerebellar functional gradients at the sub-network level. (A) Intra-cerebellar gradients. Scatterplots show intra-cerebellar gradient *I* plotted against intra-cerebellar gradients *II*-*IV* (top row), with each point representing a cerebellar vertex coloured by resting-state sub-network as defined by (Buckner et al. 2011). Positions were determined by gradient values. Ridge plots show the normalised distribution of gradient values across resting-state sub-networks for intra-cerebellar gradients *I*-*IV*. (B) Cerebro-cerebellar gradients. Scatterplots show cerebro-cerebellar gradient *I* plotted against cerebro-cerebellar gradients *II*-*IV* (top row). Ridge plots show the normalised distribution of gradient values split by resting-state sub-networks for cerebro-cerebellar gradients *I*-*IV*.

### Cerebellar functional gradients develop in the first postnatal weeks

We assessed developmental effects by aligning subject-level gradients to template-level gradients (i.e., extracted from Subset 1 consisting of the oldest subjects scanned between 43 and 44 weeks PMA) and quantifying their similarity using Spearman’s r rank correlation coefficient. Increasing PMA at scan was associated with greater subject-to-group similarity for intra-cerebellar gradient *III* (r = 0.235, p_FDR_= 0.026) and cerebro-cerebellar gradient *I* (r = 0.305, p_FDR_ < 0.001) (Figure 6), reflecting the development of macroscale organisation captured with subject-level intra-cerebellar and cerebro-cerebellar gradients.

**Figure 6.**
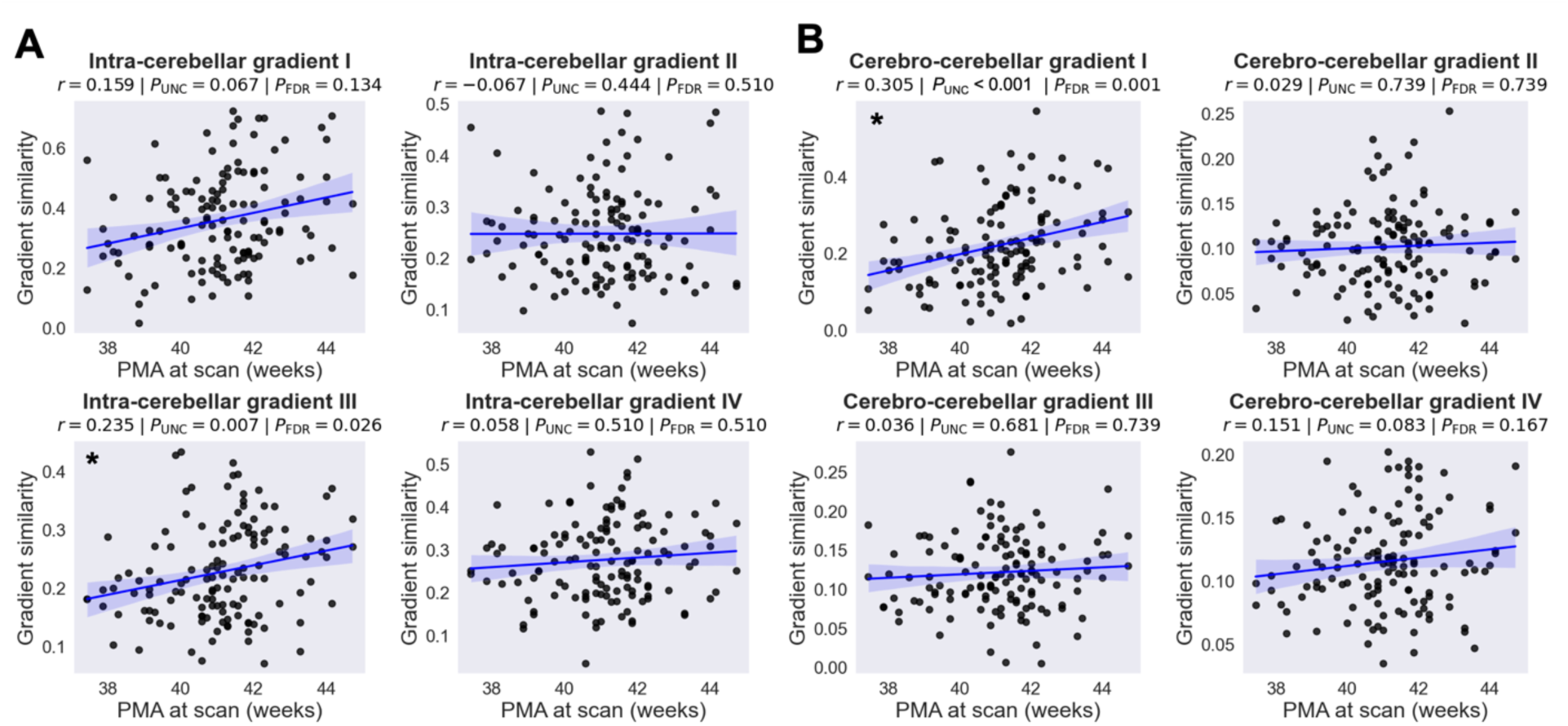
Age-related subject-to-group gradient similarity. Scatterplots showing subject-to-group template similarity plotted against PMA at scan for (A) intra-cerebellar gradients and (B) cerebro-cerebellar gradients. Each dot represents the absolute correlation coefficient between the subject and group gradient map. *p_FDR<_0.05.

## Discussion

Here we provide comprehensive mapping of functional network organisation across the human neonatal period within the cerebellum and its connectivity with the cerebrum. Using group-level ICA, voxel-wise connectivity mapping, graph theoretical metrics, and functional gradient decomposition, we show that (I) multiple ICNs within the cerebellum are already present at the time of full term gestation; (II) age-related functional changes during the neonatal period are most evident in higher-order cerebellar regions, particularly Crus I/II; (III) cerebro-cerebellar connectivity is already organised around sensory, motor and association circuits at full term gestation and strengthens with postmenstrual age; and (IV) both intra-cerebellar and cerebro-cerebellar functional gradients reflect an early continuous organisation that continues to mature across the neonatal period. Together, these findings demonstrate that neonatal cerebellar functional organisation recapitulates key principles observed in mature brains, while also undergoing dynamic and regionally heterogeneous refinement during the early postnatal period.

The early emergence of the 13 distinct cerebellar ICNs, with networks spanning most cerebellar lobules including Crus I/II as well as the vermis, highlights the early existence of functional differentiation of neonatal cerebellar organisation. The presence of networks in anatomical regions associated with both somatomotor and association functions mirrors the major canonical subdivisions and ICNs described in adults (Buckner et al., 2011; Dobromyslin et al., 2012; King et al., 2019) and suggests that the cerebellum’s foundational functional topography is already established by the time of normal birth. While previous studies have reported resting-state networks across infancy and early childhood (Lyu et al., 2025; Tikoo et al., 2025), our findings extend this work by showing substantial refinement during the narrow developmental window spanning the first postnatal weeks. In particular, the strongest age effects were found in two of the Crus I/II ICNs, with increasing coverage and connectivity strength between 37 – 44.5 weeks PMA. This finding is of particular interest as the Crus I/II region is closely tied to working memory, language, and affective processing in adults (Stoodley & Schmahmann, 2009), with its preferential connectivity with prefrontal and parietal association cortices emerging gradually across development (Bernard et al., 2016; Sydnor et al., 2021). The increasing connectivity observed in this region in the neonatal period likely reflects the earliest stages of functional specialisation within this association axis, consistent with evidence that transmodal and sensorimotor cerebellar regions of the cerebellum exhibit different patterns of development (Lyu et al., 2025). These findings parallel observations in the cerebral cortex, where large-scale somatomotor, visual, and higher-order functional networks are already evident at the time of term birth and continue to strengthen across the neonatal period (Eyre et al., 2021). Together, these results suggest that neonatal brain development is characterised by the presence of an initial functional scaffold that undergoes rapid strengthening and refinement in regions associated with somatomotor and cognitive function. Because developmental variation was modelled using postmenstrual age at scan, which captures both gestational maturation and postnatal experience, the observed age-related associations reported here likely reflect the combined influence of these processes.

Clustering the cerebrum based on its connectivity with the cerebellum revealed 15 cerebral clusters across sensory, motor, visual and associative domains. The spatial distribution of these clusters showed clear correspondence with whole-brain large-scale functional networks previously reported in neonates (Eyre et al., 2021). Cerebral visual, sensory and motor clusters showed greater alignment with such networks compared with the slightly weaker alignment of the association-related clusters, which is consistent with a developmental sensorimotor-association (S-A) axis in which cerebral somatomotor circuits consolidate early, while transmodal networks undergo more protracted maturation (Luo et al., 2024; Sang et al., 2012; Sydnor et al., 2021; Xu et al., 2026). Examination of cluster-specific cerebellar connectivity maps further revealed distinct topographic organisation within the cerebellum. Visual and association-related clusters exhibited strongest connectivity with the Crus I/II of the posterior cerebellar lobe, regions implicated in higher-order cognitive and associative processing in adult populations (Buckner, 2013; Sang et al., 2012). Motor-related clusters, predominantly connected to somatomotor-associated regions of the cerebellum including the anterior lobe and the inferior posterior lobe (Stoodley & Schmahmann, 2009), as well as the vermis which additionally receives projections from major cerebral motor networks located in primary motor and lateral and medial premotor regions (Coffman et al., 2011).

This organisation suggests that fundamental cerebro-cerebellar functional circuits are already established at birth and are arranged according to sensory, motor and association-related domains of major cerebral brain networks. This is consistent with evidence from fetal and preterm infant MRI and histology, showing that the major cerebellar input/output pathways, the inferior and superior cerebellar peduncles, are already myelinating from as early as 28 weeks gestational age (Adamsbaum et al., 2005; Brossard-Racine & Limperopoulos, 2016; Counsell et al., 2002; Yakovlev & Lecours, 1967), indicating that the structural foundation for cerebro-cerebellar communication is in place before term. Cerebro-cerebellar circuits also continue to mature rapidly across the neonatal period, reflected in increasing cluster-cerebellum connectivity strength across seven of the 15 sensory-, motor- and association-related clusters with age at scan. Together, these findings may indicate that large-scale cerebro-cerebellar networks continue to strengthen during early postnatal life. Follow-up data incorporating behavioural measures may help us determine how these early connectivity patterns relate to developing function.

We mapped functional connectivity gradients within the neonatal cerebellum for the first time, revealing that the characteristic macroscale continuous organisation seen in adults is already emerging by the time of normal birth. Both intra-cerebellar and cerebro-cerebellar gradient decompositions indicated well-defined axes of organisation, including a dominant transmodal-to-unimodal gradient analogous to the macroscale gradient architecture described in adults (Guell et al., 2018). The presence of this transmodal-unimodal axis suggests early differentiation of the regions associated with motor and higher-order functional domains, with the strong correspondence between intra-cerebellar and cerebro-cerebellar gradients suggesting that cerebro-cerebellar coupling follows shared principles of macroscale organisation from the earliest stages of development. Importantly, at the individual level, the third gradient of intra-cerebellar and principal cerebro-cerebellar functional connectivity showed increasing similarity with template gradients with age at scan, indicating that the maturation of continuous organisation principles captured by gradient analysis is detectable even across the relatively narrow early developmental window examined here. This finding supports the notion that the neonatal stage represents an important period of accelerated refinement during which the macroscale architecture of cerebellar networks begins to consolidate. Our gradient characterisation thus complements and extends the ICA and graph-theoretical results, offering a continuous, data-driven mapping of these early maturational processes.

A major strength of this study is the use of high-quality neonatal MRI from the dHCP, enabling robust mapping of cerebellar and cerebro-cerebellar organisation with unprecedented detail. Furthermore, the combination of multiple analytical approaches, including ICA, voxel-wise functional connectivity analysis, graph theory, and functional gradients, provides convergent evidence for the macroscale principles governing cerebellar network and gradient organisation at birth. However, several limitations should be acknowledged. First, although the study spans an important developmental window, this period represents only cerebellar maturation in the neonatal period. The absence of longitudinal imaging within individuals limits the ability to directly track developmental trajectories, and thus future work following infants across their first few years after birth will be essential to determine how early patterns evolve into the more differentiated organisation seen in childhood and adulthood. Second, neonatal MRI in the cerebellum remains susceptible to lower signal-to-noise ratio relative to supratentorial regions and to susceptibility-related signal dropout, which may mask more subtle connectivity and gradient maturational features.

Together, the findings presented here establish a normative framework for intra-cerebellar and cerebro-cerebellar functional organisation in the term-born neonatal brain, demonstrating the presence of intrinsic networks, continuous functional gradients, and early specialisation of both sensorimotor and association-related cerebellar regions at full term gestation. Our results further demonstrate that the subsequent weeks are a period of accelerated refinement during which cerebro-cerebellar circuits begin to consolidate their functional roles within broader neural systems. This work lays the groundwork for investigating how cerebellar circuitry develops across infancy and early childhood and provides a benchmark against which atypical development and its involvement in neurodevelopmental conditions can be assessed.

## Methods

### Participants

Subjects were selected based on qualitative and quantitative assessment of cerebellar coverage and image quality, excluding subjects with excessive in-scanner motion (>10% framewise displacement [FD] motion outliers [FD > 1mm]) and significant signal dropout in the cerebellum (> 8.5% low-signal outliers [20% below 99^th^ centile of signal intensity]). The final study sample consisted of 155 term-born neonates (67 females) from the dHCP (release 3) (Edwards et al., 2022) scanned at term-equivalent postmenstrual age (PMA) between 37.5-44.5 weeks. For most analyses, we used the 20 oldest neonates with full cerebellar coverage, which were selected as a reference sample (subset 1) and compared to the remaining 135 individual neonates (subset 2, see Supplementary Figure S1).

### MRI acquisition

We analysed fMRI data obtained as part of the dHCP at the Evelina Newborn Imaging Centre, Evelina London Children’s Hospital on a 3T Philips Achieva MRI system with a neonatal coil and imaging system (Edwards et al., 2022). Ethical approval was granted by the UK National Research Ethics Authority (14/LO/1169), and written consent was obtained from the parents or legal guardians of the infants prior to data collection. Heart rate, oxygen saturation and temperature was monitored throughout the duration of scan by a neonatal nurse or paediatrician. Blood-oxygen-level-dependent (BOLD) fMRI was acquired using a multi-slice gradient-echo EPI sequence with multiband excitation (factor 9) (repetition time (TR): 392ms, echo time (TE): 38ms, acquired spatial resolution 2.15 x 2.15 x 2.15 mm^3^ isotropic, flip angle = 34°, 45 slices, total time = 15 min 3s, number of volumes = 2300). Structural T2-weighted (T2w) images were acquired with isotropic spatial resolution 0.8 x 0.8 x 0.8 mm^3^ and a field of view of 145 x 145 x 108 mm^3^ (TR=12,000ms, TE = 156ms).

### Data processing

Individual fMRI datasets were pre-processed with the dHCP fMRI neonatal pipeline (Fitzgibbon et al., 2020). Briefly, this involved dynamic distortion and intra- and intervolume motion correction utilising a pipeline that includes slice-to-volume and rigid-body registration. FSL FIX was used to identify single-subject ICA noise components and regress out motion, cardiorespiratory, and multiband acquisition artifacts (Griffanti et al., 2014; Salimi-Khorshidi et al., 2014). The data were then registered into T2w native space using boundary-based registration (Schuh et al., 2018) and nonlinearly transformed to a 40-week neonatal template using the extended dHCP volumetric atlas. Participants showing high FD (FD > 1.5 × IQR + 75th centile) in >230 (10%) of the 2,300 volumes were excluded. The data were band pass-filtered at 0.01-0.2 Hz (Butterworth fifth order filter), smoothed with a 6mm full-width half-maximum (FWHM) Gaussian kernel, and downsampled to 2mm^3^ isotropic resolution.

### Intrinsic connectivity networks

Because of the close anatomical proximity between the cerebellum and ventral regions of the cerebral cortex, fMRI signal leakage can occur across the cerebro-cerebellar boundary, particularly near visual and anterior cerebellar motor regions. Without correction, this can result in spurious local signal overlap, leading cerebellar ICA to capture visual cortical components and reducing sensitivity to cerebellar-specific networks. To reduce influence of artefactual surrounding signal, signal from cerebral regions adjacent to the cerebellum (Supplementary Figure S6) was regressed from the 2,881 cerebellar voxel timeseries (Buckner et al., 2011). Using probabilistic group independent component analysis (GICA) on Subset 1, we decomposed the cerebellar fMRI signals into spatially distinct template ICN maps (Beckmann & Smith, 2004), also known as Resting State Networks (RSNs) (Damoiseaux et al., 2006). We used a dimensionality of 20 which allowed us to extract a number of distinct homologous unilateral and bilateral components. ICNs were named after anatomical locations in the cerebellum. For visualisation, ICN maps were projected to adult cerebellar surfaces using the SUITPy package (Diedrichsen & Zotow, 2015). Using dual regression (Nickerson et al., 2017), subject-specific spatial maps for each ICN were estimated. Core ICN amplitude was calculated as mean β-parameter value in each ICN for each subject.

### Functional connectivity analysis

dHCP specific atlases were used to extract voxel timeseries of the cerebrum, basal ganglia, thalamus, pons and cerebellum (24,838 voxels). Using Pearson’s correlation, voxel-wise functional connectivity (FC) matrices were constructed by correlating all extracted timeseries with each other in all subjects. A group FC matrix of all subjects (Subset 1) was constructed using r-to-z Fisher’s transformation.

We clustered cerebral voxels based on their correlation profiles with cerebellar voxels to characterise underlying patterns of cerebro-cerebellar connectivity, using FastICA at the group-level. We selected a dimensionality of 20, as this yielded a biologically interpretable solution comprising large-scale functional cerebral networks. Although ICA dimensionality is arbitrary and can influence the degree to which networks are split or merged, qualitatively similar patterns were observed across a range of dimensionalities. Clusters were extracted based on both quantitative and qualitative criteria. First, a cluster-size threshold was derived from a null distribution of maximum cluster extent at the selected ICA dimensionality. The cerebro-cerebellar FC matrix was permuted 1,000 times along the cerebral axis, removing spatial structure but maintaining connectivity profiles to the cerebellum. FastICA was then re-fitted to each permuted matrix using the same model order as the original analysis. For each null decomposition, the size of the largest contiguous cluster thresholded at Z ≥ 3 was recorded, yielding a distribution of maximum cluster extents expected by chance. The cluster-size threshold was set at the 95^th^ percentile of this distribution (cluster extent = 9), and only clusters exceeding this threshold were retained for further analysis. Retained clusters were visually reviewed with reference to cerebral anatomy.

At the group-level cluster degree centrality (DC) was calculated by summing connections to voxels in the resulting clusters for each cerebellar voxel, to identify region-specific patterns in the cerebellum. At the subject-level, positive connectivity values between cerebral voxels belonging to a given cluster and all cerebellar voxels were extracted and summed. This yielded a single cluster-level metric representing the total cluster-cerebellum connectivity strength. Higher values indicate stronger connectivity between the cerebral cluster and the cerebellum.

### Functional gradient analysis

Cerebro-cerebellar and intra-cerebellar FC matrices were calculated as voxel-wise Pearson’s correlation coefficients between cerebral-cerebellar and cerebellar-cerebellar time-series respectively. To reduce the impact of noise and emphasise stronger connectivitiy patterns, group and subject-level FC matrices were sparsified by retaining only the top 10% of connections in each row. Cosine similarity was then used to construct affinity matrices for all pairs of cerebro-cerebellar and cerebellar-cerebellar connectivity profiles.

Functional gradients were derived from these affinity matrices using diffusion map embedding, a nonlinear dimensionality reduction technique that identifies low-dimensional components capturing the principal patterns of variation within high-dimensional connectivity data (Guell et al., 2018; Margulies et al., 2016). Diffusion map embedding was implemented using BrainSpace (Vos de Wael et al., 2020) with the anisotropic diffusion parameter set to *α* = 0.5, a commonly used value that balances local and global relationships in the embedded space (Margulies et al., 2016; Vos de Wael et al., 2020). Gradient values place each voxel in a continuous space in which voxels with similar connectivity profiles are closely located to each other. Diffusion map embedding was first applied to the group-level cosine affinity matrices of the 20 oldest neonates (subset 1; PMA > 42.5 weeks) with the best cerebellar coverage to define neonatal cerebellar gradient templates (Oldham et al., 2025). The number of gradients was chosen based on a threshold of cumulative explained variance of 70%. Subject-level gradients were subsequently calculated for the remaining neonates (subset 2; n=135) and aligned to template gradients using Procrustes rotation (Oldham et al., 2025).

For further analysis, volume-level gradients and atlases were projected to adult cerebellar surfaces using the SUITpy toolbox (Wang et al., 2026). Gradients were contextualised in relation to cerebellar representations of adult resting-state networks, by assigning cerebellar vertices to networks using the Buckner 7-network atlas (Buckner et al., 2011) and colour-coding in scatterplots of gradient pairs accordingly. Inter-gradient similarity was assessed by correlating intra-cerebellar and cerebro-cerebellar gradients using Spearman’s rank correlation coefficient (Spearman’s *r*). Developmental effects were explored by calculating subject-to-group gradient similarity using Spearman’s *r*.

### Statistics

Summary metrics of early intra-cerebellar and cerebro-cerebellar organisation, including ICN core network strength, cerebral cluster strength, and subject-to-template gradient similarity, were associated with postmenstrual age at scan using partial Spearman’s rank correlations to examine developmental effects. Analyses were controlled for the potential confounding variables sex and total number of FD outliers. Multiple comparisons were accounted for using false discovery rate (FDR) correction (p < 0.05). The effects of PMA at scan on ICN coverage and amplitude, and cluster-cerebellum connectivity in the cerebellum were explored using general linear models (GLM) at the voxel-level, including sex and total number of FD outliers as covariates y ∼β_0_ + β_1_PMA + β_2_Sex + β_3_FD. GLM analysis was carried out using nonparametric permutation inference in FSL randomise with 10,000 permutations per test (Winkler et al., 2014). Threshold-free cluster enhancement and family-wise error rate were applied to account for potential effects of voxel-wise multiple comparisons (Smith & Nichols, 2009).

## Data and code availability

Neuroimaging and clinical data for the Developing Human Connectome Project are available via the NIMH Data Archive (collection ID: 3955; Access instructions: https://biomedia.github.io/dHCP-release-notes/).

Statistics and figures were produced in Python v3.11.9 and auxiliary packages Numpy, SciPy, nilearn, nibabel, pandas, pingouin, matplotlib, seaborn, statsmodels, SUITPy, subprocess, pathlib, os, sys, glob. Brain volume images were produced with FSLeyes. Supporting code for this manuscript is available at: https://github.com/ninatdr/cerebellum-func.

## Acknowledgments

The developing Human Connectome Project was supported by the European Research Council under the European Union’s Seventh Framework Programme (FP7/20072013)/ERC grant agreement no. 319456. This work was partly funded through the philanthropic support of Heart of Racing LLC to the Brain Health in Gen2020 programme at King’s College London, and by the National Institute for Health and Care Research (NIHR) Biomedical Research Centre (BRC): Maudsley. The authors also acknowledge support from the Institute for Translational Neurodevelopment at King’s College London, the European Autism Interventions (EU-AIMS) trial and the EU AIMS-2-TRIALS, a European Innovative Medicines Initiative Joint Undertaking under Grant Agreements No. 115300 and 777394, the resources of which are composed of financial contributions from the European Union’s Seventh Framework Programme (Grant FP7/2007–2013) and Horizon 2020 from the European Federation of Pharmaceutical industries and Associations companies’ in-kind contributions. NT, JOM, TA, DK and GM received support from the Medical Research Council Centre for Neurodevelopmental Disorders, King’s College London [MR/N026063/1]. SO was supported by The Marian E.H. Flack Trust Fellowship and the Brain and Behaviour Research Foundation (ID: 31471). QC received support from KCL-CSC joint scholarship awarded by King’s College London and Chinese Scholarship Council. DK was supported by a Wellcome Trust Career Development Award [225951/Z/22/Z]. TA was supported by a Medical Research Council (MRC) Senior Clinical Fellowship [MR/Y009665/1]. The views expressed are those of the authors and not necessarily those of the funders, IHI-JU2, King’s Health Partners, the NHS, the National Institute for Health Research, or the Department of Health and Social Care. The funders had no role in the design and conduct of the study; collection, management, analysis, and interpretation of the data; preparation, review, or approval of the manuscript; and decision to submit the manuscript for publication.

## Contributions

Conceptualisation: N.T., D.B.; methodology: N.T., S.F.M., S.O., Q.C., J.O.M., D.B.; data analysis: N.T., D.B.; interpretation of results: N.T., S.F.M., S.O., Q.C., J.O.M., D.K., T.A., G.M., D.B.; project administration: T.A., G.M.A., D.B.; supervision: G.M.A., D.B.; writing—original draft: N.T., D.B.; writing—review & editing: All authors.

