## Supplementary Information for "Early development of cerebellar functional connectivity"

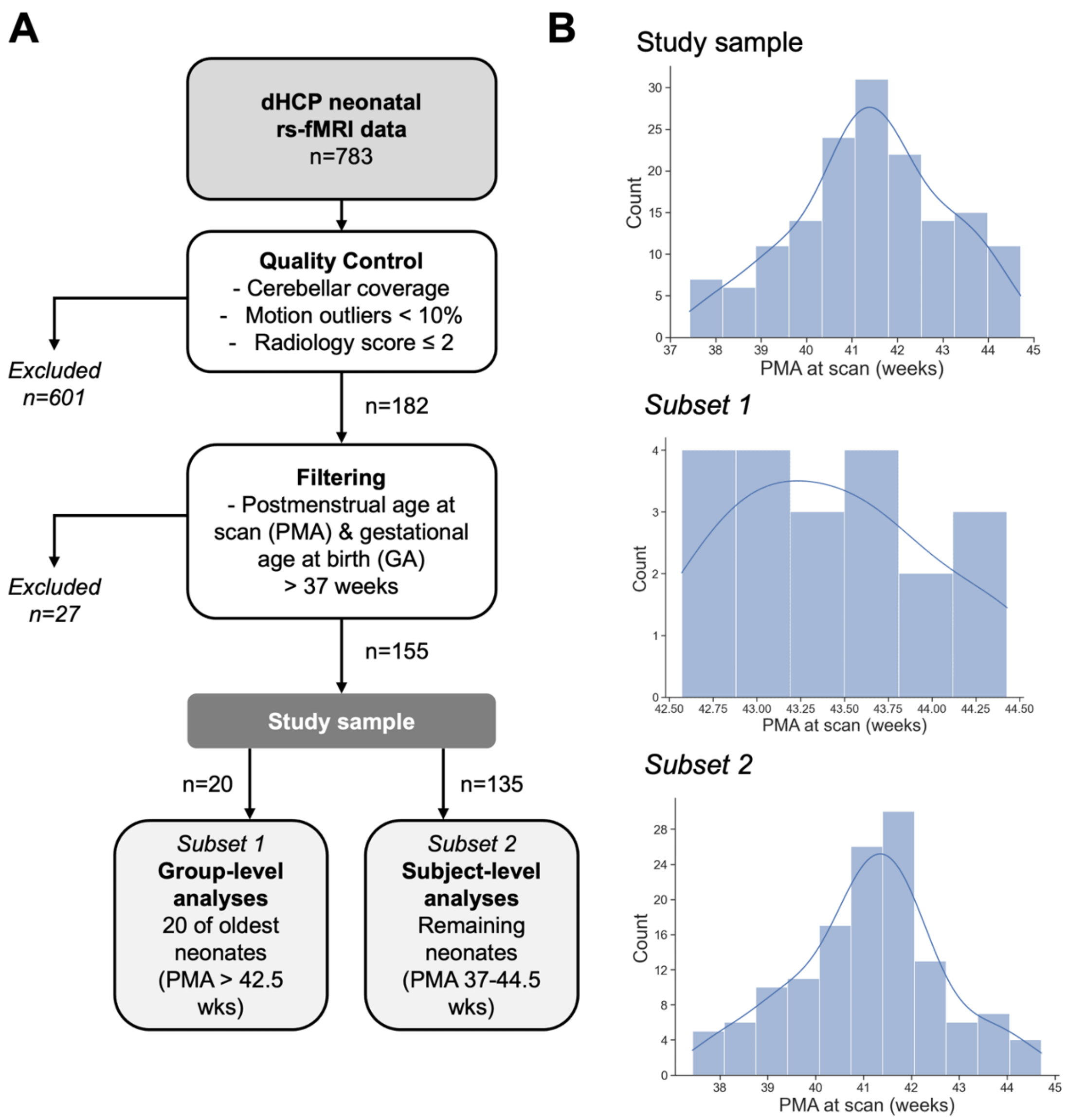


**Supplementary Figure S1.** (A) Data quality check and sample selection criteria for the total study sample and subset 1 and 2 (B) Distribution of PMA at scan (weeks) for the total study sample, subset 1 and subset 2.

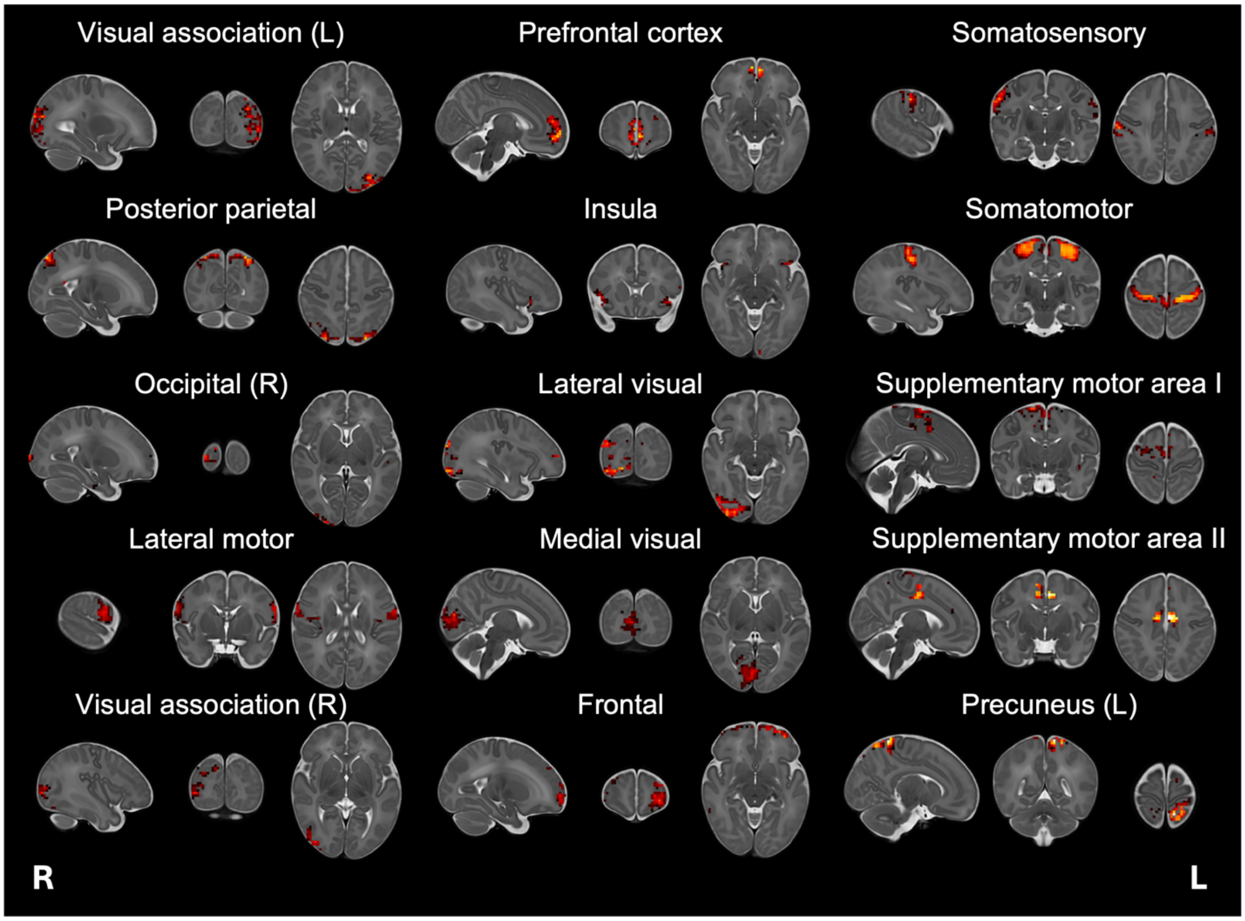


**Supplementary Figure S2.** Cerebral clusters underlying cerebro-cerebellar connectivity identified through independent component analysis (ICA). Based on permutation testing, 15 clusters were derived representing cerebral regions that share similar patterns of functional connectivity with the cerebellum. Clusters are shown in radiological orientation, neonatal template space and thresholded at $Z \geq 3$. Only components whose largest clusters exceeded permutation-derived threshold (9 voxels, 95$^{th}$ percentile of the null distribution, 1000 permutations) were retained.

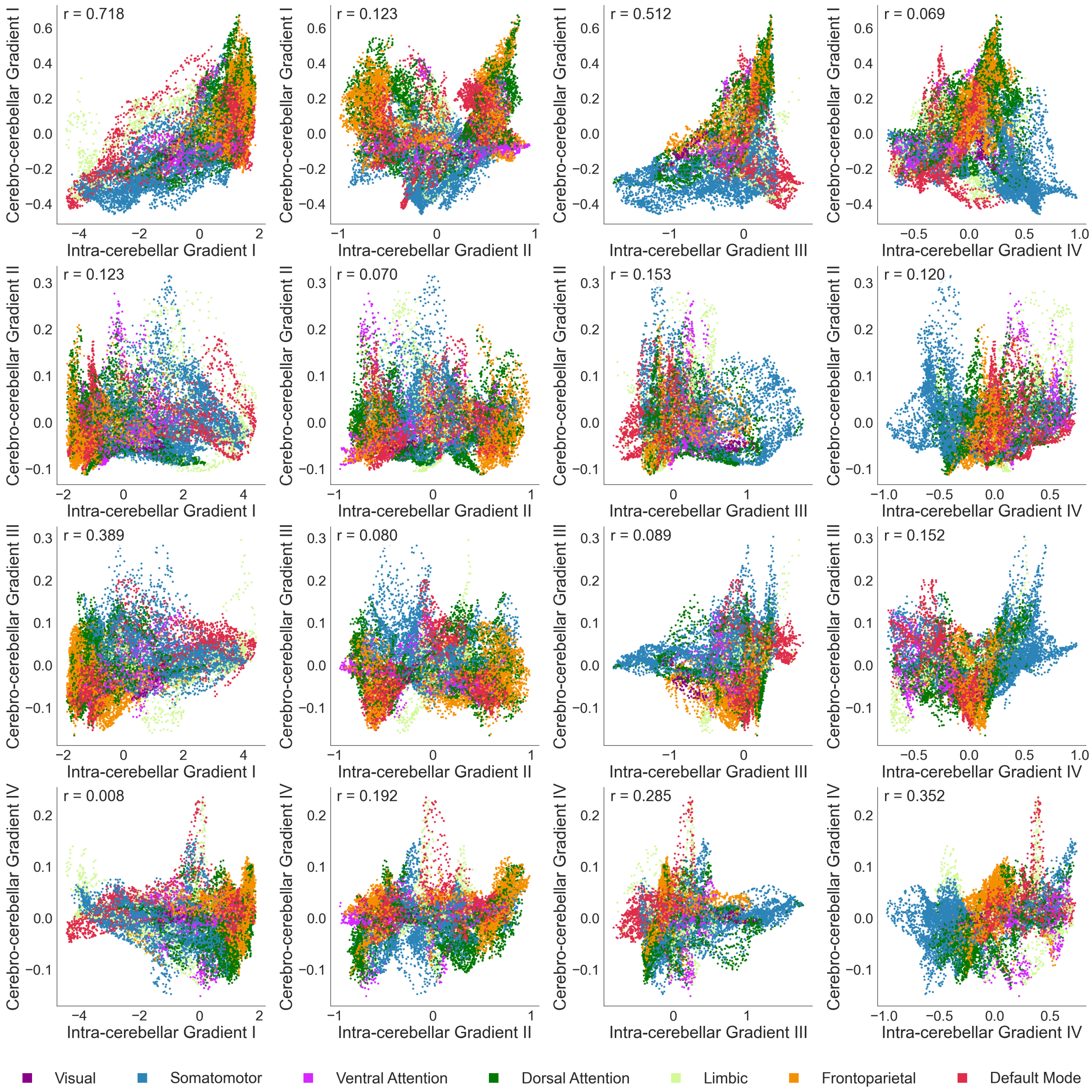


**Supplementary Figure S3.** Cerebro-cerebellar gradients I-IV (rows) plotted against intra-cerebellar gradients I-IV (columns), with each dot representing a cerebellar vertex coloured by resting-state network (Buckner et al. 2011). Positions were determined by gradient value in the respective intra-cerebellar and cerebro-cerebellar gradient space and Spearman correlations are displayed per plot.

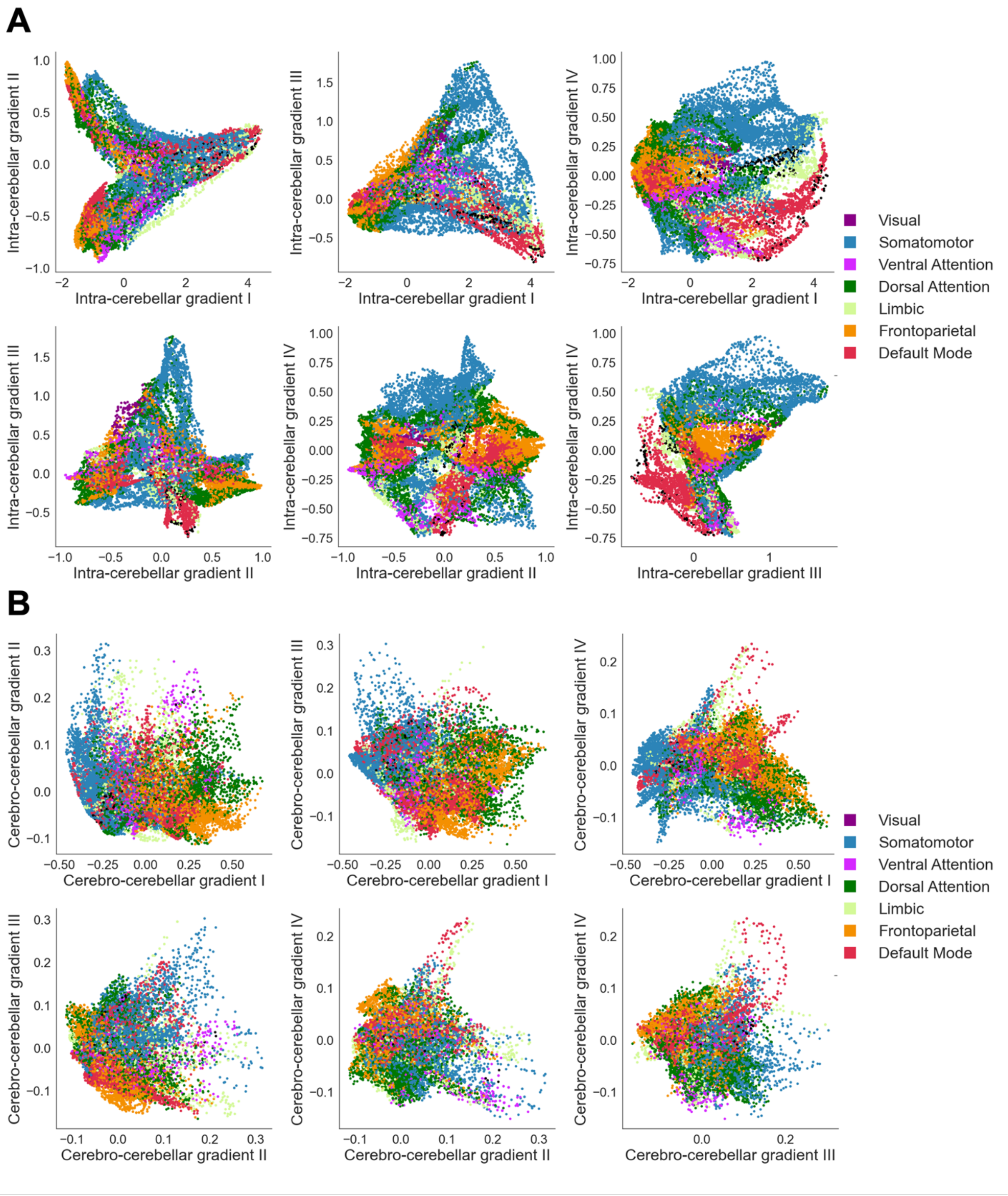


**Supplementary Figure S4.** (A) Intra-cerebellar gradients I-IV plotted against each other and (B) cerebro-cerebellar gradients I-IV plotted against each other, with each dot representing a cerebellar vertex coloured by resting-state network (Buckner et al. 2011). Positions were determined by gradient values.

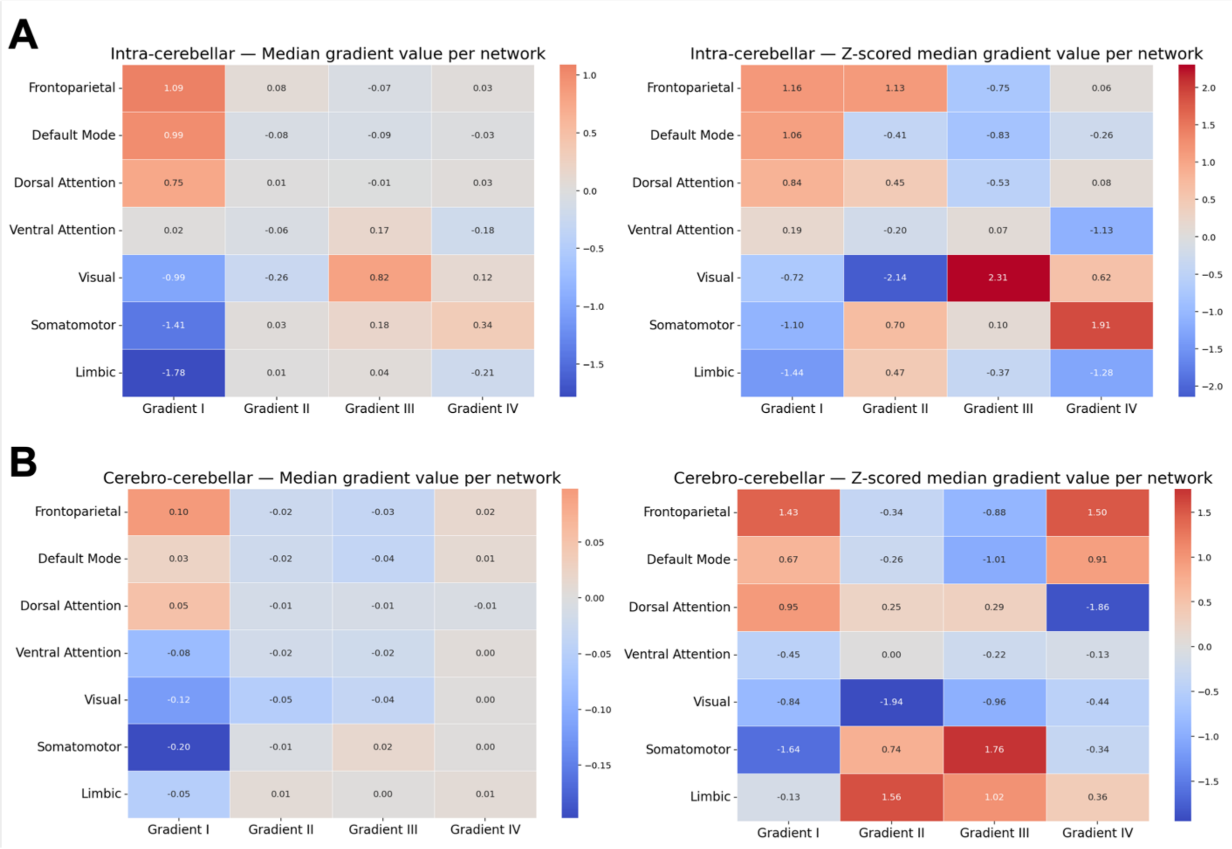


**Supplementary Figure S5** (A) Heatmap of median intra-cerebellar gradient value and (B) median cerebro-cerebellar gradient value per network for each gradient (left) and z-scored median gradient value per network (right).

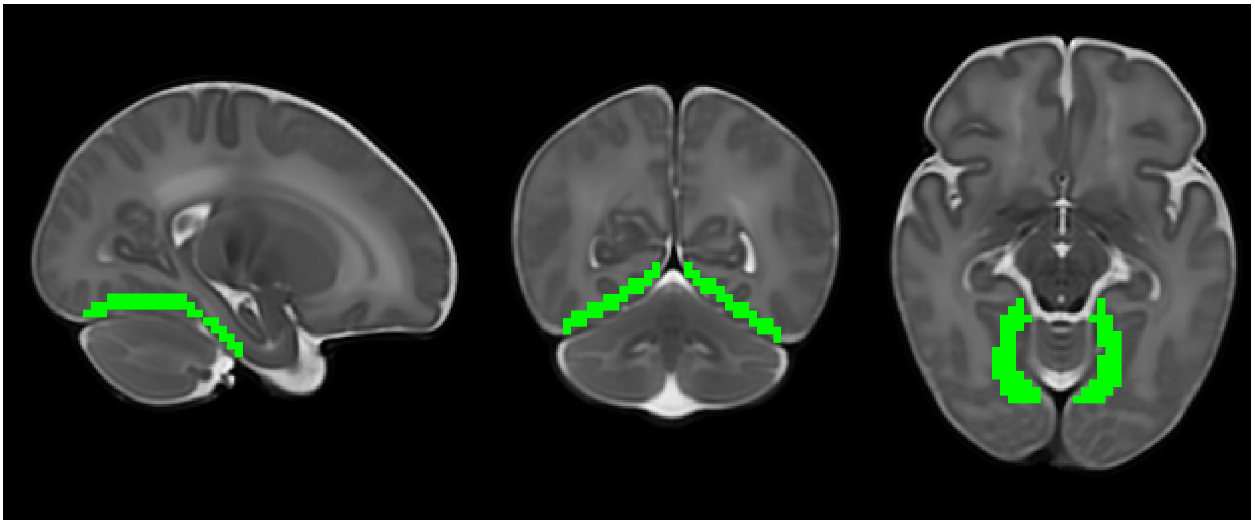

**Supplementary Figure S6.** Overlap mask in the cerebrum indicating the voxel timeseries regressed from the cerebellum before analysis.
